# *lute*: estimating the cell composition of heterogeneous tissue with varying cell sizes using gene expression

**DOI:** 10.1101/2024.04.04.588105

**Authors:** Sean K. Maden, Louise A. Huuki-Myers, Sang Ho Kwon, Leonardo Collado-Torres, Kristen R. Maynard, Stephanie C. Hicks

## Abstract

Relative cell type fraction estimates in bulk RNA-sequencing data are important to control for cell composition differences across heterogenous tissue samples. Current computational tools estimate relative RNA abundances rather than cell type proportions in tissues with varying cell sizes, leading to biased estimates. We present *lute*, a computational tool to accurately deconvolute cell types with varying sizes. Our software wraps existing deconvolution algorithms in a standardized framework. Using simulated and real datasets, we demonstrate how *lute* adjusts for differences in cell sizes to improve the accuracy of cell composition. Software is available from https://bioconductor.org/packages/lute.

## 1 Background

High-throughput bulk RNA-sequencing (RNA-seq) datasets that profile gene expression across large sample sizes are increasingly being used to identify biological differences between groups of samples, such as neurotypical control and Alzheimer’s disease cohorts (1,2). However, a major challenge with leveraging these data when profiling heterogeneous tissue is the intra-sample cell composition differences that are often observed (3). Recent efforts have been made to develop computational tools incorporating cell type-specific reference profiles based on single-cell or single-nucleus RNA-seq (sc/snRNA-seq) data to estimate the relative fractions of different cell types in bulk RNA-seq data. These estimates can be used to control for differences in cell composition across heterogenous tissue samples, which can also better determine the cell types that drive differential expression signals (4,5).

While these algorithms have been successfully used to demonstrate how cell composition changes across sample groups or conditions, an important challenge with these algorithms is that they frequently show reduced performance in heterogeneous tissues with varying cell sizes including brain (6–8), adipose (9), heart (10), and solid tumor samples (11–13). One reason for this is that the default in most deconvolution algorithms is to assume the cell sizes are the same across cell types. In this way, without adjusting for differences in cell sizes, computational algorithms estimate the relative fraction of RNA attributable to each cell type, rather than the relative fraction of cell types, leading to potentially biased estimates in cellular composition (5).

As the consequences of cell type-specific size variation started to be recognized, efforts began to incorporate cell size estimates into existing deconvolution algorithms for more accurate cell composition estimation. Improved performances after cell size adjustments were found in studies of blood (14,15) and multi-tissue (4,16) samples. The *SimBu* algorithm (17) incorporates cell size scale factors to generate bulk samples with simulated differences in cell sizes. The *ABIS* algorithm (15) uses experimentally derived and algorithmically fine-tuned cell size scale factors to improve accuracy for blood cell type predictions. The *EPIC* algorithm (12,14) adjusts on cell size prediction outputs. The *MuSiC* algorithm uses either a library normalization or user-specified cell size scaling (4). However, each of these tools were built on different frameworks with non-uniform input data formats while addressing different types of systematic errors or unwanted bias (18–25). Further, the influence of data normalizations on reference and real bulk RNA-seq is an area of active study (19). These factors can make it difficult to generate comparable deconvolution results across different algorithms, and new tools for evaluating the effects of data transformations, normalizations, and bias corrections on deconvolution outcomes are needed.

Here, we propose, *lute*, a computational tool (**Figure 1**) to accurately deconvolute cell types with varying cell sizes in heterogeneous tissue by adjusting for differences in cell sizes. The software package wraps existing deconvolution algorithms in a flexible and extensible framework to enable their easy benchmarking and comparison. For algorithms that currently do not account for variability in cell sizes, we extend these algorithms by incorporating user-specified cell scale factors that are applied as a scalar product to the cell type reference and then converted to algorithm-specific input formats. We demonstrate our method with both simulated and real experiment bulk RNA-seq data, including both heterogeneous blood (15) and brain tissues (26). While blood has been extensively studied (9,13,15,27), the brain remains mostly lacking from the literature in benchmark evaluations (26), despite the great interest and importance in determining the relative role of cell type-specific expression in heterogeneous brain tissue and their subsequent dysregulation in debilitating brain disorders (18,21). Our software is available within the Bioconductor framework (28) and can be integrated into workflows using established core Bioconductor infrastructure for bulk RNA-seq and sc/snRNA-seq data (29).

**Figure 1.**
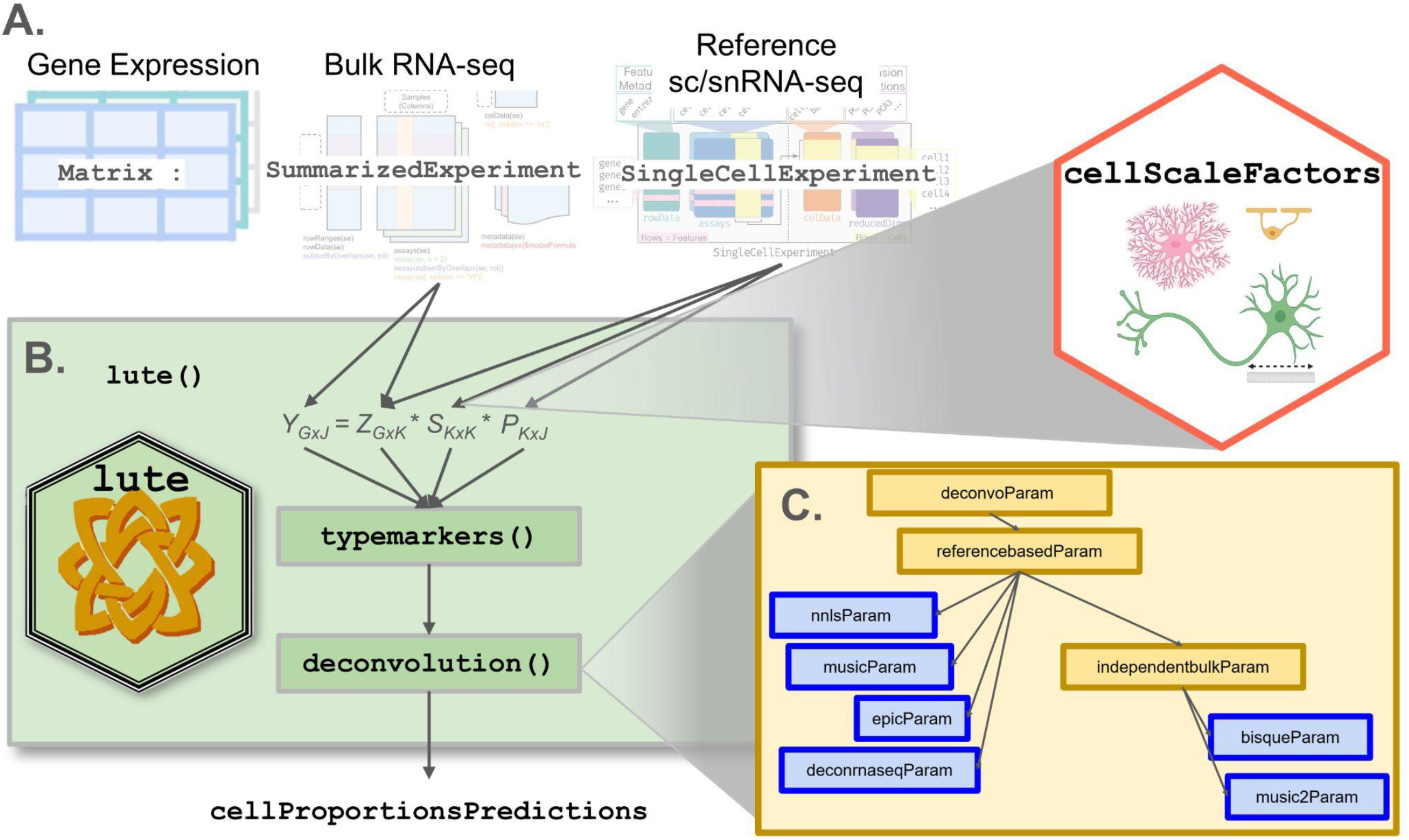
Overview of *lute* framework for deconvolution in Bioconductor. (**A**) Schematic of a deconvolution experiment using *lute*. Inputs include (far left) matrix/flat tabular format, (second from left) SummarizedExperiment (28), (second from right) SingleCellExperiment (28,29), and (far right) *cellScaleFactors*. (**B**) The *lute* framework showing (top) terms (defined in Results 2.1) for deconvolution and pseudobulk operation, including *Y* (bulk), *P* (proportions), *S* (cell type sizes), and *Z* (cell type reference), noting *k* (number of cell types) and *G* (marker genes) with arrows indicating applicable input classes and scaling factors corresponding to differences in cell sizes across cell types available in the *cellScaleFactors* (hexagon) R/Bioconductor package (18), (middle) the `typemarkers()` function to select marker genes from the reference and bulk, and (bottom) the `deconvolution()` generic for calling the user’s choice of the deconvolution method. (**C**) Schematic illustrating supported deconvolution algorithms in *lute* including a parent class (a.k.a. “deconvoParam”), referencebasedParam, independentbulkParam, nnlsParam, musicParam, epicParam, deconrnaseqParam, bisqueParam, and music2Param.

## 2 Results

### 2.1 *lute*: deconvolution of heterogeneous tissue with varying cell sizes

We begin with a general formulation of cell type deconvolution to demonstrate how to adjust for differences in cell sizes as introduced previously (18), followed by a summary of the *lute* software package. Consider a set of high-dimensional *Y*_*GxJ*_ representing a heterogeneous tissue sample from *g* ∈ (1,…, *G*) marker genes expressed and *j* ∈ (1,…, *J*) bulk RNA-sequencing samples. We assume the heterogeneous tissue is a mixture of *K* cell types indexed by *k* ∈ (1,…, *K*). Using a referenced-based sc/snRNA-seq approach, the standard equation to estimate the cell composition of *J* bulk RNA-seq samples is *Y*_*GxJ*_ = *Z*_*GxK*_ * *P*_*KxJ*_ where the goal is to estimate *P*_*KxJ*_ the *K* cell type proportions each of the each of the *J* bulk samples using a cell type-specific reference matrix *Z*_*GxK*_ containing for *G* marker genes across the *K* cell types.

Next, consider a vector of scalars *s*_*K*_ = (*s*_1_,…, *s*_*K*_) representing the cell size for each *k* ^*th*^ cell type, which could be computationally derived or, most often, experimentally derived from an external dataset, ideally from an adjacent tissue slice (18). We can define the matrix *S*_*K*_ = *I*_*KxK*_ * *s*_*K*_ where *I* _*KxK*_ is an identity matrix. Then, in a similar manner as above, if we consider the equation *Y*_*GxJ*_ = *Z*_*GxK*_ * *S*_*KxK*_ * *P*_*KxJ*_, we can define a new matrix *Z*′_*GxK*_ = *Z*_*GxK*_ * *S*_*KxK*_ and solve for *P*_*KxJ*_ using an equation similar as above *Y*_*GxJ*_ = *Z*′_*GxK*_ * *P*_*KxJ*_ (Methods). In this way, we estimate the cell composition *P*_*KxJ*_ while also adjusting for differences in cell size. We note that, without scaling by *S*_*KxK*_, the assumption is that cell sizes are equal. For example, in *lute*, the default algorithm is non-negative least squares (*NNLS*) (30) where for each *j*^*th*^ sample,*p*_*kj*_ > 0 and 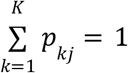 and user-specified cell scale factors *s*_*K*_ are applied as a scalar product to the cell type reference and mapped to inputs for the deconvolution algorithm. However, *lute* supports *NNLS (30), MUSiC (4), MuSiC2 (16), EPIC (14),DeconRNASeq (10)*, and *Bisque (9)*.

To address the problem of independent deconvolution frameworks with non-uniform input data formats, we were inspired by the *bluster* Bioconductor package (31) designed to address a similar problem for unsupervised clustering algorithms. We take standard Bioconductor S4 classes as input, including SummarizedExperiment (32), SingleCellExperiment (28,32), and a vector of cell sizes, either user-provided or loaded from the *cellScaleFactors* (33) ExperimentData package (**Figure 1A**). Then, we define a S4 generic function called `deconvolution()` and create separate S4 classes in a hierarchy for each algorithm supported (**Figure 1B-C**). This facilitates modular support for algorithms available across multiple repositories, including CRAN (9,30), Bioconductor (10), and GitHub (4,14,16). For example, deconvolution algorithms that depend on the existence of reference-based sc/snRNA-seq profiles all share a common S4 class (34) (**Methods**). In this way, for each algorithm, *lute* is able to map standard data inputs *Y, Z*, and *S* (also described in **Box 1**) to the appropriate algorithm-specific synonyms and implementations.

#### Box 1

**Summary of key terms**. Column 1 refers to the terminology introduced in Figure 1 and used throughout the manuscript. Column 2 gives the definition of the term.

| Terminology | Definition |
| --- | --- |
| sample | tissue slice (from a tissue block) or cell mixture from tissue |
| donor | subject-level identifier for source of tissue |
| bulk sample | tissue sample containing mixture of cells across $K$ cell types ($Y$) |
| cell type reference matrix | matrix containing cell type-specific gene expression for $G$ genes across $K$ cell types ($Z$) |
| cell size factors | vector of cell sizes of $K$ cell types ($S$) |
| cell composition | vector of the proportion of $K$ cell types in a given bulk sample ($P$) |
| RMSE | square root of the mean of squared errors |

### 2.2 Application of *lute* using simulated pseudobulk data

In the next two sections, we considered two applications of *lute* using *in silico* pseudobulk data, where we simulated bulk RNA-seq profiles by aggregating sc/snRNA-seq data mixed together in various known proportions or cell compositions. We demonstrate how differences in cell sizes lead to inaccurate estimates of cell composition, but scaling for differences in cell size leads to improved accuracy for cell composition estimation. In addition, there is great interest in investigating how cell composition changes in the brain, particularly the human dorsolateral prefrontal cortex (DLPFC), are associated with neurodegenerative and neuropsychiatric disorders including Alzheimer’s Disease (AD) (7), major depressive disorder (35), and schizophrenia (36,37). Recent evidence in schizophrenia suggested gene expression changes accompany onset (36,38,39), while other studies showed neuroinflammation, mediated by non-neuronal cells called microglia, is linked to early stages of neuropsychosis (40). High cellular heterogeneity in DLPFC makes deconvolution challenging, partially and a known component of this is due to structural and microenvironmental complexity arising from six distinct cortical layers (41). Similar independent analyses showed there are many molecularly distinct subpopulations among the 6 previously mentioned fundamental brain cell types of inhibitory neurons, excitatory neurons, oligodendrocytes, oligodendrocyte precursor cells, astrocytes, and endothelial cells (42). Specifically, we considered plasmablasts compared to other blood and immune cell types from peripheral blood mononuclear cells (PBMC) isolated from whole blood, where cell size adjustments showed improvement in bulk transcriptomics deconvolution accuracy (12,14,15). Plasmablasts, otherwise known as antibody-secreting cells, have distinct transcriptional activity from other blood cell types and are studied for their roles in febrile vasculitis (43) and autoimmunity (44,45). For both of these tissues (brain tissue and blood tissue), we demonstrated how differences in cell sizes lead to inaccurate estimates of cell composition using *in silico* pseudobulk and real bulk RNA-seq samples.

#### 2.2.1 Example from human postmortem DLPFC

We demonstrate the performance of *lute* by simulating pseudobulk RNA-seq data using a snRNA-seq dataset (46) from neurotypical postmortem human DLPFC brain tissue with cell types that we aggregate to *k*=2 cell types, namely neurons (excitatory and inhibitory) and glia (oligodendrocytes, oligodendrocyte precursor cells, astrocytes, and microglia). Briefly, we show performance improvement in estimating the cell compositions with and without adjusting for cell sizes (**Figure 2A**). In this brain region, it is known that neurons are nearly 3x larger than glia (47), which makes it an illustrative dataset to demonstrate the performance improvements from cell size scale factor normalization with *lute*. In this dataset, we utilized *N*=17 snRNA-seq libraries generated from tissue blocks obtained from 10 adult neurotypical donors across three regions of the DLPFC including the anterior, posterior, and mid regions. The snRNA-seq from each tissue block had a median of 3,004 nuclei per sample (**Table S1**) and cells were aggregated to create 17 pseudobulk profiles with neuron-glia ratio ranging between 80/20% to 25/75% cell composition. Pseudobulks were generated using the product of a specified (or known) cell type proportions and reference snRNA-seq mean expression profiles. Cell size rescaling was performed by taking the scalar product of cell type expression and a set of cell size scale factors (**Table S2, Methods**). We performed feature selection using the snRNA-seq data to identify the top 40 cell type marker genes using the mean ratio of the sample-adjusted expression (*Methods 4*.*2 Marker selection*) (26). Using these markers, we used *NNLS* (30) to estimate the cell composition of the pseudobulk samples for *k*=2 groups.

**Figure 2.**
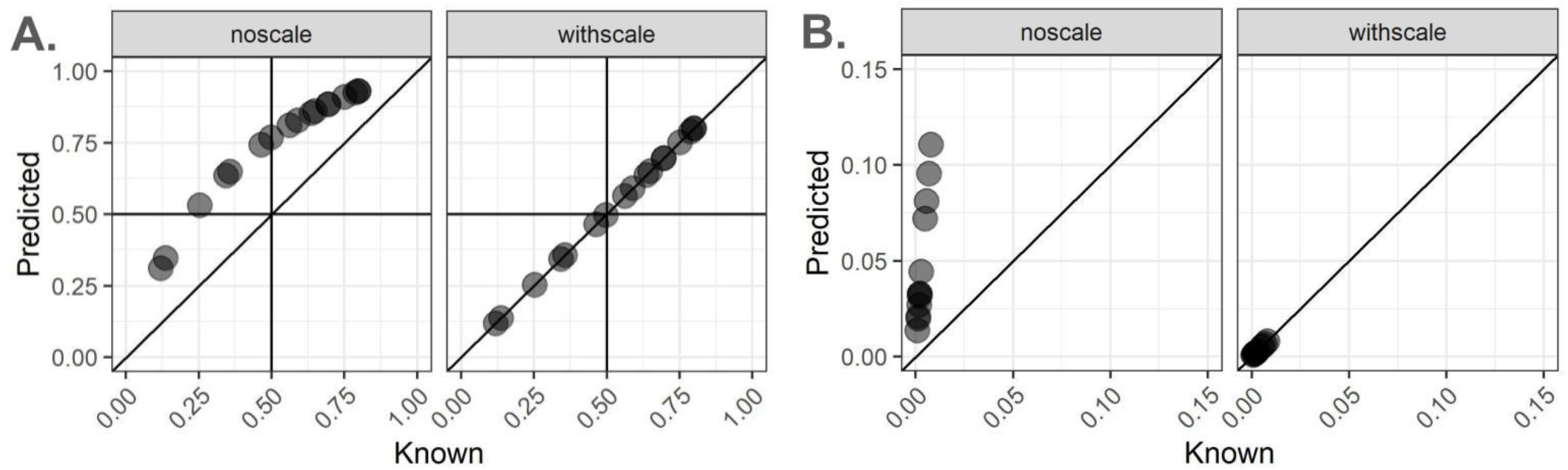
Performance improvement when adjusting for differences in cell sizes with *lute*. Pseudobulk samples were created using sc/snRNA-seq data and mixing cell types together with a known cell type proportion (*x*-axis). The predicted cell type proportions (*y*-axis) were estimated using *NNLS* with and without adjusting for differences in cell sizes. The known (*x*-axis) and predicted (*y*-axis) cell type proportions are shown without scaling (left) and with scaling (right). (**A**) *N*=17 pseudobulk profiles were created by mixing neuron and glia cell types at a prespecified ratio ranging between 80/20% to 25/75% cell composition using *N*=17 snRNA-seq libraries generated from tissue blocks obtained from 10 adult neurotypical donors in Huuki-Myers et al. (2023) (46) across three regions of the dorsolateral prefrontal cortex (DLPFC). (**B**) *N*=12 pseudobulk profiles created mixing plasmablasts-other cell types ranging between 7.04*10^-4^ - 1.47*10^-3^ % to 0.992 - 0.999 % cell composition using bulk RNA-seq based reference profile of peripheral blood mononuclear cells (PBMC) (15). Diagonal lines indicate y = x and no error.

Without accounting for differences in cell sizes in *lute* between neurons and glia, we observed an overestimation of the proportion of neurons and an underestimation of the proportion of glia cells (root mean squared error, RMSE = 0.22) (**Figure 2A, Table S3**). This overestimation reflects the algorithm estimating the relative fraction of RNA attributable to each cell type, rather than the relative fraction of cell types, leading to biased estimates in cell composition (5). However, when adjusting for differences in cell sizes (**Table S2, Methods**), we found more accurate estimates of cell composition (RMSE not scaling = 0.22, RMSE scaling = 1.34*10^-16^) (**Figure 2A, Figure S1A-B**). We found similar results (RMSE not scaling = 0.17, RMSE scaling = 9.38*10^-17^) when expanding to *k*=3 cell types (excitatory neurons, inhibitory neurons, and glia) using the same dataset (**Figure S2A-B, Table S3**). In this analysis, expanding from *k*=2 to *k*=3 had only a slight impact on error in glial cell estimates (not scaling, RMSE_k2_ - RMSE_k3_ = 0.22 - 0.17 = 0.04), which diverged from prior findings in brain tissue (48).

To assess the robustness of *lute* to other reference profiles, we repeated the pseudobulking experiments with a different snRNA-seq dataset from postmortem human neurotypical DLPFC (49). Here, the snRNA-seq data were from *N*=3 donors with a median of 4,209 nuclei per sample (**Table S4**), which were aggregated to create pseudobulk profiles with 19-60% neuron and 12-48% glia cell composition and were adjusted for differences in cell sizes (**Table S5**). Using the same feature selection as above, we estimated the cell composition for *k*=2 (**Figure S1C-D**) and *k*=3 (**Figure S2C-D**) groups of cell types and found similar results to the first DLPFC dataset (**Figure 2A, Table S3**).

Finally, we investigated whether the scaling factors used to adjust for differences in cell sizes that were uniquely derived for each dataset could be generalized to adjust for differences in cell sizes in other datasets. Specifically, using the snRNA-seq libraries generated from DLPFC tissue blocks obtained from 10 adult neurotypical donors, we estimated the cell sizes for cell types within each snRNA-seq sample using marker library expression and paired orthogonal in situ hybridization (smFISH) measurements (**Methods, Table S6**). Next, we randomly shuffled the smFISH cell sizes (**Table S7**) across the *N*=13 snRNA-seq libraries derived from unique DLPFC tissue blocks (cell sizes neuron, mean = 37.04, median = 36.09, sd = 4.29; cell sizes glial mean = 30.51, median = 30.59, sd = 2.33). The purpose of this is to simulate the scenario where we are interested in estimating the cell composition from a bulk RNA-seq tissue sample, but we have not directly measured the cell sizes for each cell type. We asked whether cell sizes measured using smFISH/immunofluorescence (IF) data using RNAScope/IF technology in one snRNA-seq sample can be generalized to other samples. We found that randomly shuffling the cell size across tissue blocks used as input to *NNLS* did not lead to a reduction in performance (**Table S7**), showing that estimating the cell composition is robust when using cell sizes calculated from a different source (**Figure S3, Table S8**). While RMSE reduction from this experiment was consistent across donors (RMSE noscale = 0.05, withscale = 0.03, **Table S3**), this improvement was non-uniform because of differences among cell scale factors used to generate the pseudobulk (i.e. taken as the matrix product of reference and cell size estimates) and the cell scale factors used to perform the deconvolution. This demonstrated that analysis with *lute* facilitates fine tuning of cell scale factors under differing simulation conditions, including cell scale factor and marker expression magnitudes.

#### 2.2.2 Example from human PBMC

Next, we considered a different tissue with cell types that also vary size, namely peripheral blood mononuclear cells (PBMC). The dataset profiled 29 immune and blood cell types from *N*=12 healthy young adult donors for which known (a.k.a. “true”) cell proportion estimates were available, and it was used to calculate the ABsolute Immune Signal (ABIS) PBMC cell type reference. Reference profiles from these data were based on purified bulk RNA-seq transcripts per million (TPM) expression, and known estimates of cell composition came from flow cytometry cell abundances. PBMC features heterogeneity in cell sizes (27), and plasmablasts are known to be larger than other cell types in this tissue by up to 15.32 fold (15). The reference contained *N*=4 donors with a median of 2.0*10^6^ cells per sample (50), which were aggregated to create pseudobulk profiles with 9.00*10^-4^ - 0.008% plasmablasts and 0.992 - 0.999% other cell composition. Using the same feature selection as above, we estimated the cell composition for *k*=2 and found improvement (RMSE unscaled = 5.37*10^-02^, scaled = 6.63*10^-17^ **Table S3**) for the estimates of cell composition for plasmablasts (**Figure 2B, Figure S4**).

### 2.3 Application of *lute* using observed bulk RNA-seq data

In this section, we used real (or observed, not *in silico* pseudobulk) bulk RNA-sequencing data to evaluate accuracy of the deconvolution algorithms used to estimate the cell composition of heterogeneous tissue with varying cell sizes. Using the data described in Section 2.2.1, there were a subset of *N*=12 DLPFC tissue blocks that had paired bulk RNA-seq along with the snRNA-seq data along with matched smFISH/IF data (**Table S8**) (26). We found that adjusting for differences in cell sizes using *NNLS* led to an improved performance in terms of estimating the cell composition (**Figure 3**). We also compared the performance of *NNLS* to (i) *MuSiC (4)*, as it uses gene variance-based scaling to improve across-sample integration in multiple samples, and (ii) *Bisque (9)* (**Figure S5**), as it adjusts for assay-specific biases and was shown to outperformed other algorithms in recent analyses of human cortex (6). No differences were observed from *Bisque* with or without rescaling (**Figure S5C-D**), reflecting the fact that this algorithm’s linear adjustment method effectively adjusts away the effect of taking the scalar product of cell size scale factors. With known neuron proportions calculated as fraction total cells from snRNA-seq, correlations (**Figure S6**, Pearson’s R coefficient) were highest in either scaling condition for *Bisque* (both R = 0.76), followed by *NNLS* with cell size scaling (R = 0.65), *MuSiC* with scaling (R = 0.63), and *NNLS* (R = 0.33) and *MuSiC* (R = 0.30) without scaling. One outlier sample (sample id: Br8667_mid) showed high unscaled RMSE (*NNLS* = 0.48, **Figure 3A-B**, *MuSiC* = 0.48, **Figure S5A-B**) that was reduced by similar magnitude after scaling (*NNLS* R = 0.41, *MuSiC* R = 0.42) *MuSiC* and *NNLS*. This sample showed the lowest error from *Bisque* (R = 0.15). In summary, while *Bisque* showed the best performance before normalization, *NNLS* and *MuSiC* tied for best performance after cell size scale factor normalization. However, the ideas in *lute* to adjust for cell sizes can easily be integrated into any deconvolution algorithm.

**Figure 3.**
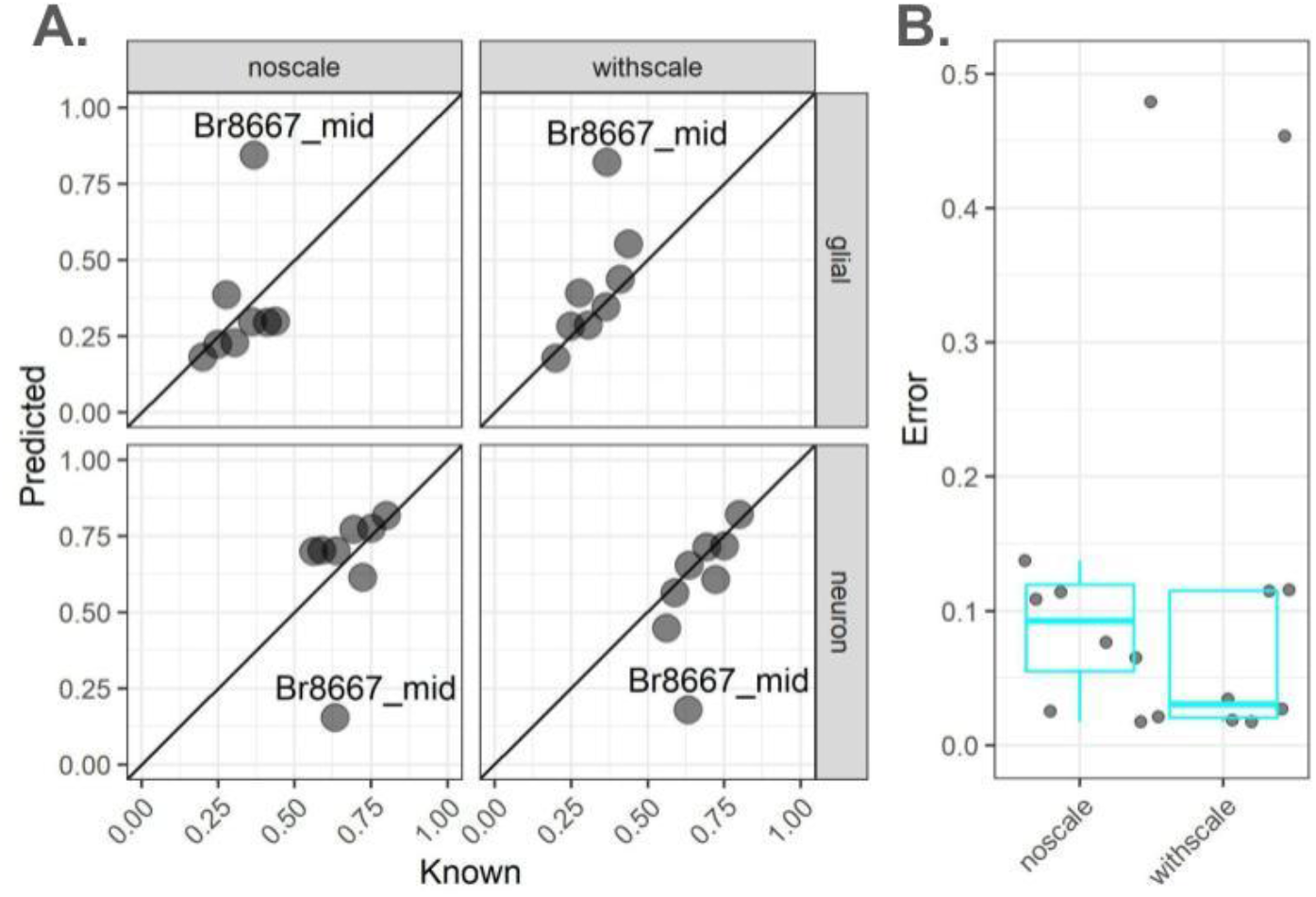
Estimates of the proportion of neurons in observed bulk RNA-seq DLPFC samples using *NNLS*. Analysis of *N*=8 observed bulk RNA-seq DLPFC samples from neurotypical donors from (26). (**A**) Scatterplots of (x-axis) known versus (y-axis) predicted neuron and glial cell proportions using *NNLS* without scaling (left column) or with cell size factor scaling (right column) for bulk RNA-seq samples from DLPFC in which known cell type proportions are estimated from snRNA-seq data. Text label indicates Br8667_mid, an outlier sample. Diagonal lines indicate y = x and no error. (**B**) Jittered points and quantile boxplots showing (y-axis) error either (left) with or (right) without scaling.

## 3 Discussion and conclusions

In this paper, we introduce a software package *lute* that can be used to accurately estimate the proportion of cell types with varying cell sizes in heterogeneous tissue by adjusting for differences in cell sizes. Our package *lute* wraps existing deconvolution algorithms in a flexible and extensible framework to enable easy benchmarking and comparison of existing deconvolution algorithms. We performed a comparison of three algorithms, *NNLS, Bisque*, and *MuSiC*, and found cell scale factor adjustment improved outcomes for these algorithms. This indicates cell scale factor adjustment could be more useful in settings where cell type heterogeneity is of greater concern (i.e. in specific experiments or tissues), or where available samples are not matched by donor (i.e. where sample sources are discordant).

While benchmark evaluations have emerged for deconvolution, large RNA-seq datasets featuring matched orthogonal measures suitable for systematic deconvolution experiments are lacking for brain and other tissues because it is difficult to systematically profile all cell types within them (20). Where few samples and nuclei are available per study, multiple datasets can be used to make the cell type reference for deconvolution (4,7,51), and our findings showed scaling on cell size scale factors should work in these settings. Furthermore, we replicated prior findings (5) that cell types having high mRNA size scale factor bias show systematic over-prediction from deconvolution, and that cell types having very low bias show systematic under-prediction. Here we fill in the gaps of existing benchmark studies and demonstrate applications of *lute* to experimental and simulated bulk RNA-seq data from heterogenous brain and blood tissues.

While we performed normalization as an explicit and discrete step upstream and independent of downstream algorithmic deconvolution, this opens the door for further experimentation to either fine-tune normalizations or show algorithm performance either with or without cell size scaling. Importantly, *lute* supports such investigations, which flexibly allows a user-defined marker gene identification algorithm and deconvolution algorithm prior to cell size normalization.

Marker gene quality and efficacy is another open topic for bulk transcriptomics deconvolution of heterogeneous tissues, likely due to several factors. Recent methods have mapped canonical cell type expression markers onto private or difficult-to-access datasets (22,23). However, it remains uncertain whether a canonical or private cell type marker expression reference is preferable for deconvolution. Also, no consensus standard exists for marker gene selection, and multiple available methods have not been formally tested for deconvolution of heterogeneous tissues.

In conclusion, *lute* allows characterization of consistent deconvolution improvements from normalization, and these improvements are a function of the selected cell type markers and known size in real and simulated data. Our package is available on Bioconductor and can be used to extend and improve existing deconvolution algorithms by adjusting for differences in cell sizes. We aim to encourage researchers to embrace cell size rescaling as a new standard processing step to develop and test bulk transcriptomics deconvolution techniques, which will expedite further breakthroughs in the transcriptomics field.

## 4 Methods

### 4.1 Data

#### 4.1.1 Overview of paired bulk-RNA-seq, snRNA-seq and smFISH/IF datasets from adult neurotypical postmortem human DLPFC tissue blocks

As previously described by Huuki-Myers et al. 2023 (46), paired (meaning both types of -omics were measured on the same *N*=19 tissue blocks) snRNA-seq measured on the 10x Genomics Chromium platform and spatially-resolved transcriptomics data measured on the 10x Genomics Visium platform were generated from tissue blocks collected from 3 different positions across the anterior-posterior axis of the DLPFC (designated posterior, anterior, and middle) from 10 adult neurotypical control donors (**Table S1**). In our analyses here, we only use the snRNA-seq libraries to generate the pseudobulk RNA-seq profiles. In addition to the snRNA-seq libraries generated from these tissue blocks, there was also paired bulk RNA-sequencing, and single molecule fluorescent in situ hybridization (smFISH)/immunofluorescence (IF) using RNAScope/IF technology generated and described in Huuki-Myers et al. 2024 (26) measured in the same tissue blocks. The smFISH/IF data was used to measure the cell type composition in the same tissue samples serving as a “gold standard” to compare the estimated cell composition in the bulk RNA-seq predicted by the deconvolution algorithms in (26). The fact that all three (snRNA-seq, bulk RNA-seq, and smFISH/IF) technologies were measured on the same tissue blocks helps to minimize potential donor-specific unwanted variation or batch effects.

##### 4.1.1.1 Preprocessing of snRNA-seq and smFISH/IF data from DLPFC tissue blocks

Out of the *N*=19 tissue blocks from Huuki-Myers et al. 2023 (46), *N*=13 tissue blocks (excluding samples Br2720_post and Br6471_ant) had matched snRNA-seq and smFISH/IF data at the time of analysis. We used the *N*=17 snRNA-seq libraries measured on the 10x Genomics Chromium platform from these 10 adult neurotypical donors obtained from across three regions of the dorsolateral prefrontal cortex (DLPFC) (**Table S8**). These snRNA-seq libraries were used to create the *in silico* pseudobulk RNA-seq profiles (**Figure 2A**), and the subset of N=13 pseudobulk samples with matched smFISH/IF data were used to perform shuffle experiments (**Figure S3**).

The preprocessing and initial cell type label assignment of these snRNA-seq data were described previously (26). Nuclei with outlying high mitochondrial gene expression and low gene expression, consistent with run failure, were removed. Cell type labels were initially mapped to snRNA-seq data using a multi-step clustering strategy. We removed cells not labeled as neuronal or glial from this strategy (e.g. immune cells, etc.) prior to downstream analyses. We identified six distinct cell types (labeled “Inhib” for inhibitory neurons, “Excit” for excitatory neurons, “Oligo” for oligodendrocytes, “Astro” for astrocytes, and “EndoMural” for Endothelial and Mural cells), which we resolved into *k*=3 (Inhib, Excit, glial) and *k*=2 (neuron and glial) label sets. We combined these cell types under the broad labels of “glial” (“Oligo”, “Astro”, “EndoMural”) and “neuron” (“Excit” and “Inhib”. For pseudobulking experiments we implemented two cell type label resolutions of combined as k2 (“neuron” and “glial”) and k3 (“glial”, “Excit”, and “Inhib”, **Table S2**).

For the smFISH/IF data, fluorescent labels were developed for the RNAScope/IF assay and imaged using the HALO software. Two RNAScope/IF probe combinations marked 3 cell types each: the first (*N=*12) included Excit, Micro, and Oligo/OPC; the second (*N=*13) included Astro, Endo, and Inhib (see Huuki-Myers et al. 2024 (26) for further details on smFISH/IF preprocessing). In the analyses here, the cell types labels from smFISH/IF and snRNA-seq were approved by consensus from three image analysis experts.

##### 4.1.1.2 Preprocessing of bulk RNA-seq data from DLPFC tissue blocks

From the same 10 DLPFC donors, paired bulk-RNAseq data was collected from 19 tissue blocks using three different RNA extraction methods: (i) isolated total RNA, (ii) isolated nuclear RNA, and (iii) isolated cytosolic RNA. Two RNA-seq libraries were prepared from each RNA sample using either RiboZero Gold or PolyA library preparation techniques. Bulk RNA-seq processing and quality control is described in Huuki et al. 2024 (26), a total of 110 bulk RNA-seq samples were produced by this dataset, with a maximum of 6 per tissue block. For the purposes of our analyses, we did not distinguish between the different RNA extraction methods or RNA library types. Of *N*=19 the DLPFC tissue blocks, our analyses used a subset of *N*=8 tissue blocks from *N=6* donors that had smFISH data matched with bulk RNA-seq from nuclei prepared using RiboZeroGold (Results Section 2.3, **Figure 3**).

In addition to the above, all bulk RNA-seq samples passed further additional minimum quality filters, with a minimum of 38,750 (median = 704,918) counts marker expression, and a maximum of 30 (median = 1) zero-expression markers, by sample (Supplementary Materials). Two tissue blocks had two samples of matched bulk RNA-seq and snRNA-seq data (Br8667 middle and anterior; Br8492 middle and posterior). In assessments of cell amount accuracy across conditions, we calculated RPKM and applied a log transformation using pseudocounts with log2 normalization (logNormCounts function from the `scuttle` (v1.12.0) (52) R/Bioconductor package).

##### 4.1.1.3 Estimating cell sizes using RNAScope/IF data

To estimate cell sizes using smFISH/IF, cells were imaged and processed in HALO (Indica Labs). A maximum nucleus area of 78µm was applied to remove out-of-focus cells. Image expected nuclei count was filtered according to a maximum nuclei count of 1,362,399, which was determined using a quantile filter of 97%. Cell sizes were calculated from RNAScope/IF as the median for each of six broad cell type markers detected using HALO. RNAScope/IF data labeled 6 broad cell types across the DLPFC: excitatory neurons, inhibitory neurons, oligodendrocytes, astrocytes, microglia, and endothelial cells.

RNAScope confidence filters were defined by expert image analyst review (26). Images were processed as pairs for each tissue slice, and pairs were graded at three quality levels. After filtering to retain the two highest quality levels, we used 12 samples with image-based cell sizes from at least 1 slice including 9 samples with both paired slices in downstream analyses.

#### 4.1.2 Overview of snRNA-seq from adult neurotypical postmortem human DLPFC tissue blocks in Tran et al. (2021)

Next, we used *N*=3 DLPFC snRNA-seq libraries from Tran et al. (2021) generated from DLPFC from 3 adult neurotypical donors. These snRNA-seq samples were used in the *in silico* pseudobulk experiments in this paper. The snRNA-seq libraries were generated using the 10x Genomics Chromium platform. The preprocessing for the snRNA-seq data was described previously(49).

#### 4.1.3 Overview of bulk RNA-seq from PBMC samples

Bulk RNA-seq data were processed as described in (15). We further performed simulations using previously published median transcripts per million (TPM) of bulk RNA-seq data from purified cells (a.k.a. the *ABIS* reference) and flow cytometry cell abundances from PBMCs of 12 healthy individuals in total (15). Gene names were mapped to Ensembl IDs using the *biomaRt* (v2.58.0) (53) R/Bioconductor package. Cell types were binarized as either “Plasmablast” or “Non-plasmablast” combining 16 cell types, including, MAIT, NK, and multiple types of dendritic cells, Monocytes, naive T-cells, and memory T-cells. After removing cell types absent in the reference, flow cytometry (a.k.a. “known”) proportions ranged from 7.01*10^-4^ - 0.776 for plasmablasts and 0.001-0.276 for non-plasmablasts. In mathematical terms defined previously, *Z* was from bulk RNA-seq data from 4 donors and *P*_*known*_ was based on flow cytometry data from 12 donors. After generating (*Y* = *Z* * *S* * *P*_*known*_) 12 pseudobulk samples, NNLS was used to obtain 12 *P*_*predicted*_ vectors for the two (K = 2) cell types of interest (**Figure S7**).

### 4.2 Marker selection on bias-adjusted expression

We used a two-step pipeline to adjust snRNA-seq data for batch effects. Adjustments were performed separately for each cell type categorization scheme (e.g one adjustment series each for K2 and K3, respectively). First, we adjusted for sample batch effects using `ComBat()` from the *sva* (54) (v3.44.0) R/Bioconductor package. We ran this function in parametric mode and specified the cell type labels as the principal covariate. In the second step, we used the *scuttle* (v1.6.3) (52) R/Bioconductor package to downsample counts according to minimum library size observed across batches within each cell type.

Cell type gene marker selection from snRNA-seq data was performed on the batch-adjusted normalized log-transformed expression. We identified the most reliable cell type markers at three resolutions as the markers with the highest concordance (i.e. occurring as markers for the same cell type consistently across all slides) and overlap (i.e. occurring in at least 3 of 12 slides) across sample sources. Markers were identified using the *Mean Ratio* of cell type expression with the `get_mean_ratios2()` from the *DeconvoBuddies* (v.0.99.0) R package (26). At 80 marker genes per cell type selected using the highest ratio of mean expression, this resulted in a median of 286 counts per cell, and a median of 1 zero-expression markers, by cell (**Table S1**).

### 4.3 Deconvolution algorithms with *NNLS, MuSiC*, and *Bisque*

We used our *lute* (v.0.99.30) R/Bioconductor package to perform deconvolution using several algorithms. *NNLS* was accessed using a *lute*-compatible class wrapper to call the `nnls()` (30) function from the *NNLS* (v1.4) R package. *MuSiC* (4) was accessed using a *lute*-compatible class wrapper, which called the `music.basic` function from the *MuSiC* (v.1.0.0) (55) R package from GitHub. *Bisque* (9) was accessed using the `ReferenceBasedDecomposition` function from the *BisqueRNA* (v.1.0.5) (56) R/CRAN package. Class wrappers for deconvolution algorithms are described in the *lute* companion vignette on Bioconductor. We performed experiments with and without rescaling before deconvolution with either *NNLS, MuSiC*, or *Bisque*.

### 4.4 Cell size scale factors

Cell size scale factors used in rescaling were computationally and experimentally derived (**Table S2, Table S5, Table S6**) (26). Experimentally derived factors were calculated based on high-resolution image capture from RNAScope/IF experiments followed by processing with the HALO (v3.3.2541.383, Indica Labs) image analysis software. Cells were labeled with DAPI, a nuclear marker, and cell type-specific fluorescence markers, as well as a fluorescent marker for *AKT3*, a size-specific marker across cell types (47). We further summed expression marker counts for each cell type prior to conducting experimental bulk RNA-seq analyses (**Table S2**). We selected manual cell size scale factor integers of 10 for neuron and 3 for glial (ratio = 3.33) that fell between orthogonal scale factor ratios for median marker expression (*k*=2, neuron/glial = 11.58) and previously published scale factors for these cell types (5).

### 4.5 Pseudobulking experiments

To better understand bias due to differences in cell sizes, we performed a pseudobulking experiment series across samples and cells passing quality filters from multiple human DLPFC cohorts (46,49), where cell type labels were assigned by the same clustering pipeline in each cohort. We used the *lute* function `ypb_from_sce()` to generate pseudobulk samples as the matrix product of the proportions and cell type reference. For example, in pseudobulking the DLPFC samples, we manually set a large divergence in cell sizes between all neuron and glial labels, where neurons, including Inhib an Excit, had a manually set scale factor size of 10 and glial, including Oligo, Astro, Micro, and EndoMural, had a size of 3, and the scalar product was then taken with the cell reference atlas using *lute* (**Figure S3**). Rather than simulate cell proportion mixtures, we confined study to empirical reference-proportion combinations to demonstrate real snRNA-seq dataset utility, as samples containing low cell type proportions may have distinct expression patterns at markers compared to high cell type proportions (48).

Simulations testing the impact of cell size factors on deconvolution outcomes in terms of bias and RMSE used the following mathematical approach. First, we defined the generative function for a simulated bulk sample such that the matrix product was computable:

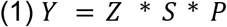

Where *Y* is a matrix of *G* markers by *J* samples, *P* is a vector of proportions of length equal to *K* cell types, *S* is a vector of *M* scale factors of length *K*, and *Z* is the signature matrix of dimensions *G* markers by *K* cell types.

Next, suppose we compare two formulations for *Z*:

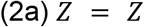

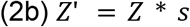

The first formulation (2a) is the marker gene expression summarized across cells for each of the *K* types, without additional rescaling or adjustment. The second formulation (2b) is equivalent to (2a) after rescaling by taking the scalar product of the *S* cell size factor vector. Finally, we obtain the following estimates for *P*:

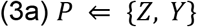

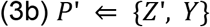

In (3a) and (3b), we use the same function “⇐” to obtain two sets of estimated cell type proportions. These are *P* based on the unscaled signature matrix *Z*, and *P*′ based on the rescaled signature matrix *Z*′.

### 4.6 Performance metrics

Error was calculated as the absolute difference between known and predicted proportions.

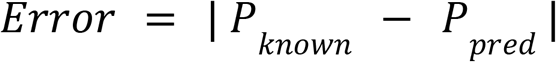

To assess the accuracy, we used root mean squared errors (RMSE) across cell types:

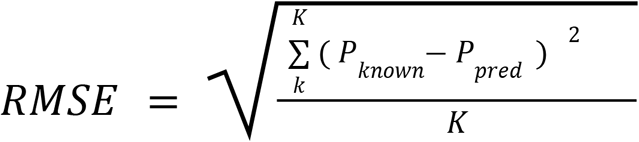

Where *K* is the total number of cell types, *k* is the *k*th cell type, *P*_*known*_ is the known (a.k.a. “true”) cell type proportion in the *k*th cell type, and *P*_*pred*_ is the predicted cell type proportion in the *k*th cell type. RMSE calculations were identical in cohorts 1 and 2, and calculations for *k*=3 included three cell types (inhibitory neurons, excitatory neurons, and glial), with a further calculation of *k*=3 neuron as the sum of predictions in inhibitory and excitatory neurons.

### 4.7 Shuffling pseudobulk experiment factors

Suppose we adapt the pseudobulk equation from (1) as follows:

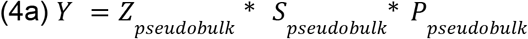

Then from (3a) we have

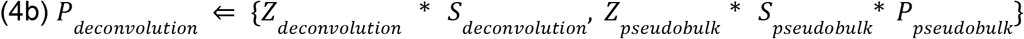

Where *S*_*deconvolution*_ has dimensions identical to *Z*_*deconvolution*_. We designate terms separately for *pseudobulk* and *deconvolution* with subscripts. We performed shuffling experiments for each of the five terms in (4b), in which one term was held constant while the others were either matched or from one of the remaining samples in (46), and we repeated this experiment for a low- and high-neuron sample.

### 4.8 Statistical analyses

Statistical analyses used base R (v4.2.2) packages and functions. Simulations and random sampling were performed using the base R functions `rnbinom()` for the negative binomial distribution, `rnorm()` for the normal distribution, `rpois()` for the poisson distribution, and `sample()` for random vector selection. All operations incorporating randomizations were initiated using `set.seed()` for computational reproducibility. Plots were generated using *ggplot2* (v3.3.6) and *GGally* (v2.1.2).

## Declarations

### Ethics approval and consent to participate

Not applicable.

### Consent for publication

Not applicable.

### Competing interests

The authors declare that they have no competing interests.

### Availability of data and materials

We provide access to the integrative multi-assay human DLPFC dataset, and interactive notebooks and scripts to reproduce our analyses at https://github.com/LieberInstitute/deconvo_lute-paper (50). The *lute* software package is available from GitHub and Bioconductor (https://bioconductor.org/packages/lute).

### Funding

This project was supported by the Lieber Institute for Brain Development, and National Institutes of Health grant R01 MH123183. All funding bodies had no role in the design of the study and collection, analysis, and interpretation of data and in writing the manuscript.

### Author contributions

SKM and SCH wrote the initial draft and edited the manuscript. SKM and SHK prepared the figures. SKM prepared the tables. LAHM, LCT, and KM contributed to the conceptualization of the manuscript and provided comments on the draft. All authors approved the final manuscript.

## Acknowledgements

We would like to thank Kelsey D. Montgomery and Sophia Cinquemani (LIBD) for contributing to the collection of imaging data analyzed in this manuscript. While an Investigator at LIBD, Andrew E. Jaffe helped secure funding for this work. Schematic illustrations in **Figure 1** were generated using BioRender.

## Supplemental figures

**Figure S1.**
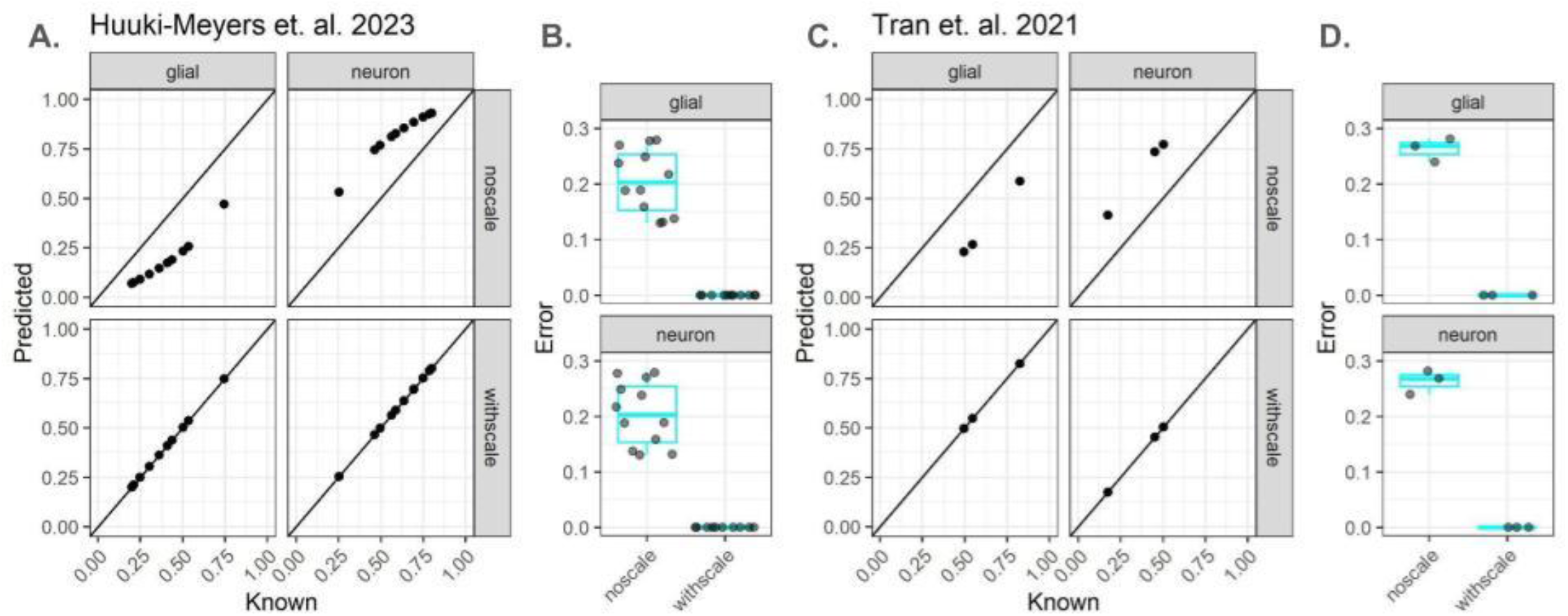
Pseudobulk simulation results from two DLPFC datasets with *k=2* cell types. (**A-B**) Pseudobulk results from Huuki-Myers et al. (2023) (46). (**A**) We estimated the cell composition for *k*=2 (neuron and glia) resolution using *NNLS* without (top row) and with (bottom row) scaling for differences in cell sizes, where the known cell composition is on the *x*-axis and the estimated cell composition is on the *y*-axis. The figure is faceted by cell types (neuron and glia) along the columns. (**B**) Boxplots of the absolute error (magnitude difference between the known and predicted cell composition) for the *N*=12 pseudobulk samples, for (top) glia and (bottom) neuron. (**C-D**) Pseudobulk results from Tran et al. (2021) (49). (**C**) We estimated the cell composition using *k*=2 (neuron and glia) using *NNLS* without (top) and with (bottom) scaling for differences in cell sizes where the known cell composition is on the *x*-axis and the estimated cell composition is on the *y*-axis. The figure is faceted by cell types (neuron and glia) along the columns. (**D**) Boxplot of the error (difference between the known and predicted cell composition) for the *N*=3 pseudobulk samples, for (top) glia and (bottom) neuron. Diagonal lines indicate y = x and no error.

**Figure S2.**
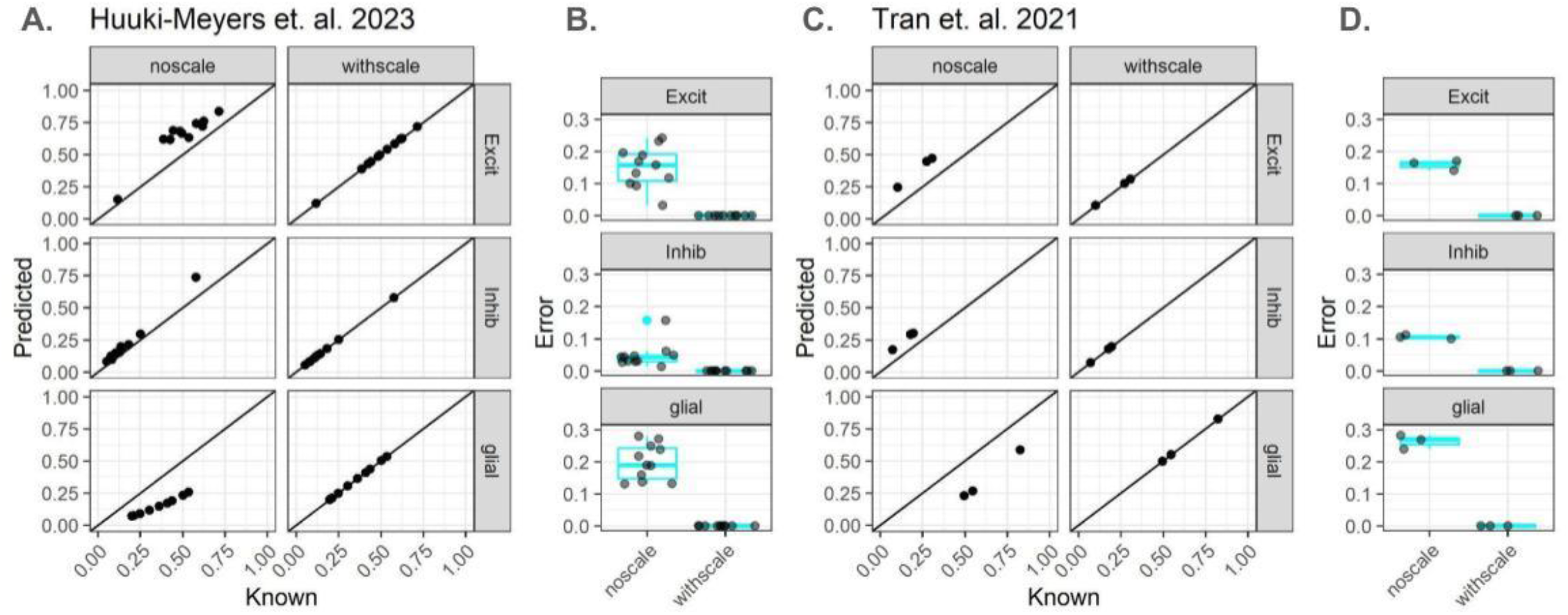
Pseudobulk simulation results from DLPFC datasets with *k=3* cell types. (**A-B**) Pseudobulk results from Huuki-Myers et al. (46). (**A**) Scatterplots of estimated the cell composition using *k*=3 (excitatory neuron, inhibitory neuron, and glia) using *NNLS* without (left) and with (right) scaling for differences in cell sizes; the known cell composition is on the *x*-axis and the estimated cell composition is on the *y*-axis. The figure is faceted by cell types (neuron and glia) along the rows. (**B**) Boxplots of the error (difference between the known and predicted cell composition) for the *N*=12 pseudobulk samples. (**C-D**) Pseudobulk results from Tran et al. (2021) (49). (**C**) Scatterplots of the cell composition using *k*=3 (excitatory neuron, inhibitory neuron, and glia) using *NNLS* without (left) and with (right) scaling for differences in cell sizes where the known cell composition is on the *x*-axis and the estimated cell composition is on the *y*-axis. The figure is faceted by cell types (neuron and glia) along the rows. (**D**) Boxplots of the error (difference between the known and predicted cell composition) for the *N*=12 pseudobulk samples. Diagonal lines indicate y = x and no error.

**Figure S3.**
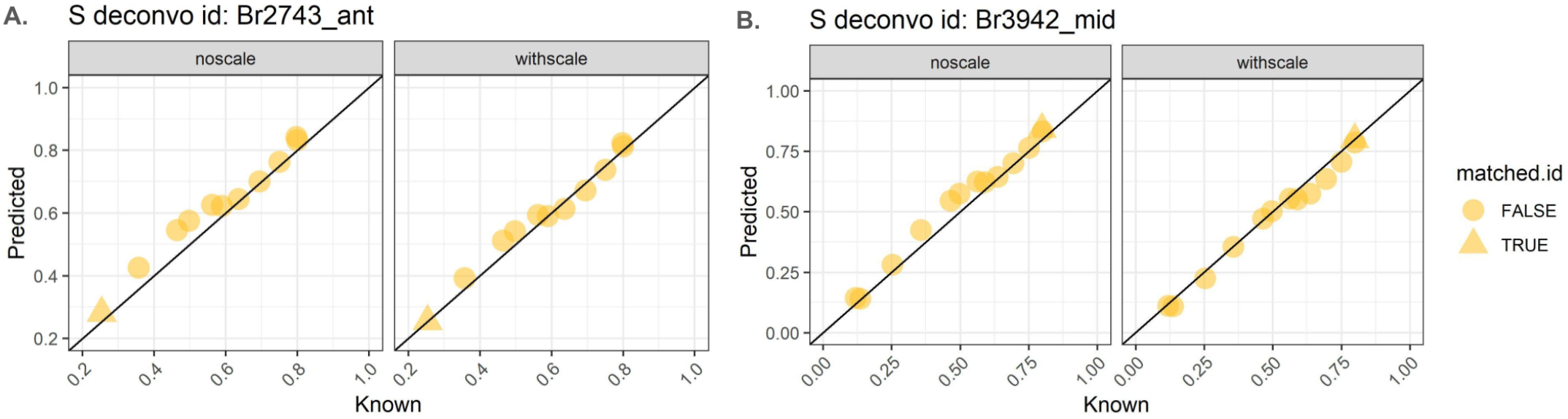
Impact of randomly shuffling RNAScope cell scale factors in pseudobulk simulations. (**A**) Scatterplots of either (left panel) with or (right panel) without adjusting by cell sizes from sample with low neuron proportions (plot title, Br2743_ant). (**B**) Scatterplots of either (left panel) with or (right panel) without adjusting by cell sizes from sample with high neuron proportions (plot title, Br3942_mid, **Table S7**). Points correspond to if the cell sizes were matched (triangle) or unmatched (circle), where references and cell scale factor arrays were calculated from DLPFC dataset (47). Diagonal lines indicate y = x and no error.

**Figure S4.**
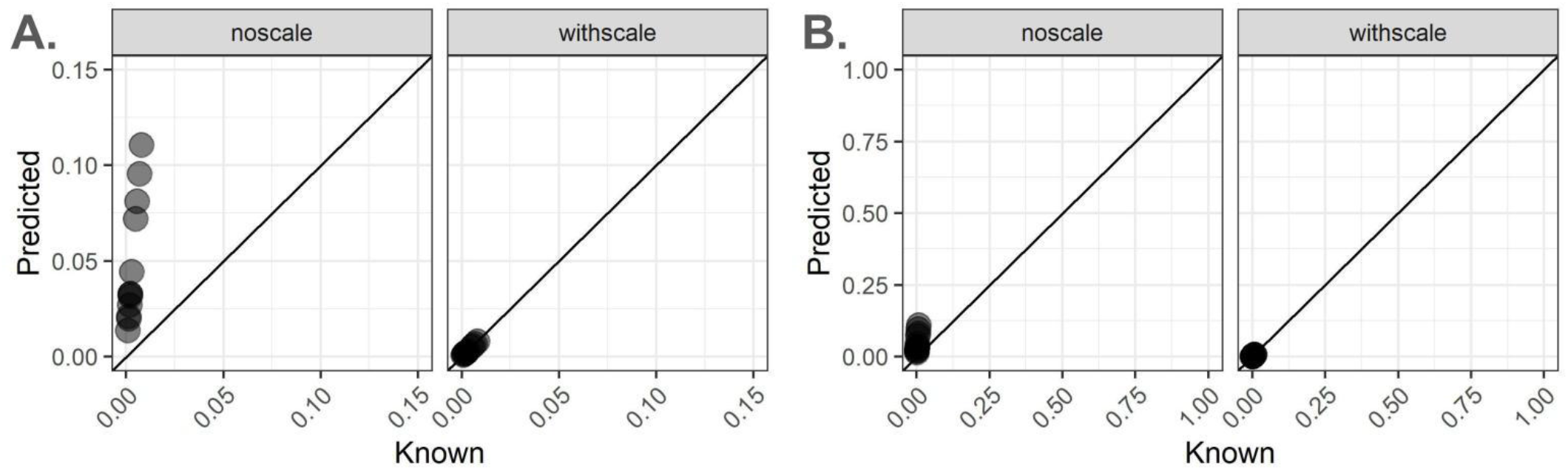
Deconvolution results before and after rescaling in an independent PBMC experiment. Results are shown for *N*=12 samples from (15) with known proportions from flow cytometry (**Methods**) at two zoom levels, either (**A**) axis maximum = 0.15 or (**B**) axis maximum = 1. Scatterplots show the (x-axis) known flow cytometry proportions versus the (y-axis) predicted proportions of Plasmablasts either (right panel) before or (left panel) after rescaling on cell size scale factors. Diagonal lines indicate y = x and no error.

**Figure S5.**
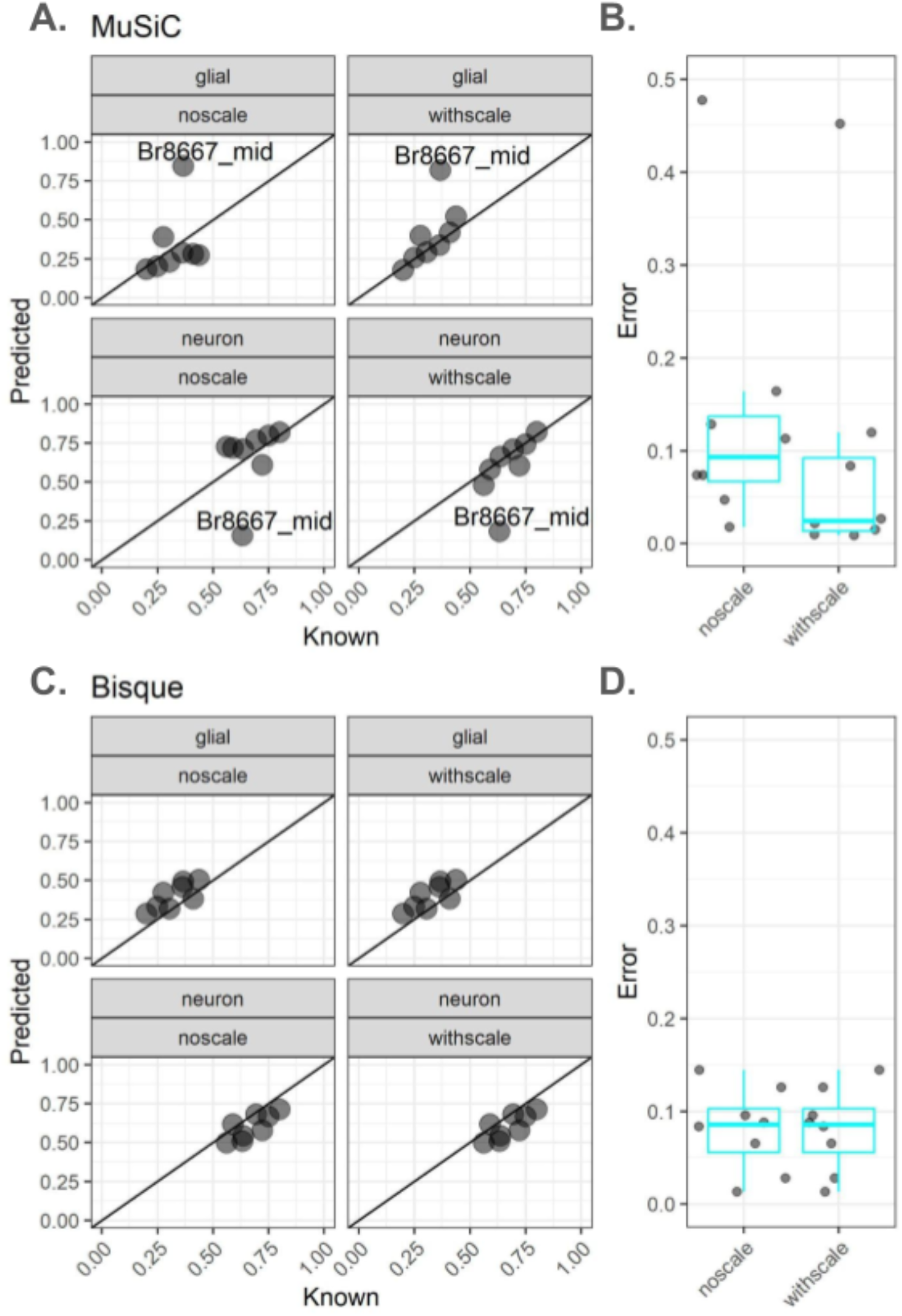
Results of neuron predictions across deconvolution algorithms in experimental DLPFC RNA-seq samples from (46). (**A**) Scatterplots show results from *MuSiC* in (points) experimental DLPFC bulk RNA-seq samples (top row) glial and (bottom row) neurons either (left column, “noscale”) without scaling or (right column, “withscale”) with scaling, with text label indicating outlying sample. Diagonal lines indicate y = x and no error. (**B**) Jittered points and quantile boxplots of (y-axis) errors by (x-axis) scaling. (**C**) Scatterplots show results from *Bisque* in (points) real bulk RNA-seq samples (top row) glial and (bottom row) neurons either (left column, “noscale”) without scaling or (right column, “withscale”) with scaling. Diagonal lines indicate y = x and no error. (**D**) Jittered points and quantile boxplots of (y-axis) errors by (x-axis) scaling.

**Figure S6.**
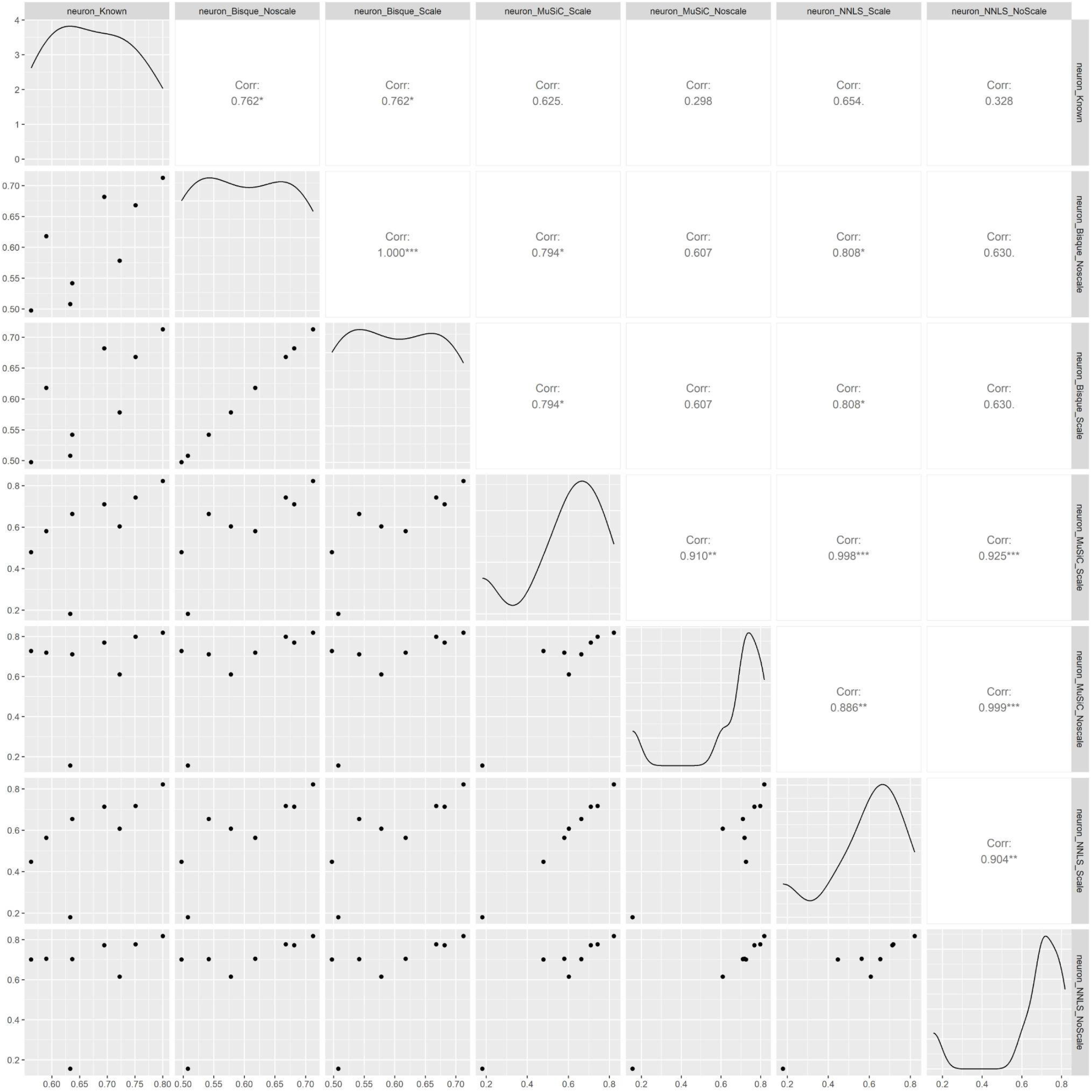
Correlations of predicted and known neurons in experimental DLPFC bulk RNA-seq, from (47), for *NNLS, MuSiC*, and *Bisque*. Pairs plots generated using `GGally` of known and predicted neuron proportions from multiple algorithms in real bulk RNA-seq samples from multiple preparation conditions, for neuron. Row and column labels indicate the cell type, algorithm (either “nnls” for *NNLS* (30), “music” for *MuSiC* (55), or “bisque” for *Bisque* (9), or known), and condition (either scale or noscale). Text panels contain the Pearson R correlation magnitude, with asterisks indicating significance (none : 0.10 <= p; . : 0.05 < p < 0.10; * : 0.01 < p < 0.05; ** : 1.0*10^-3^ < p < 0.01; *** : p < 1.0*10^-3^).

**Figure S7.**
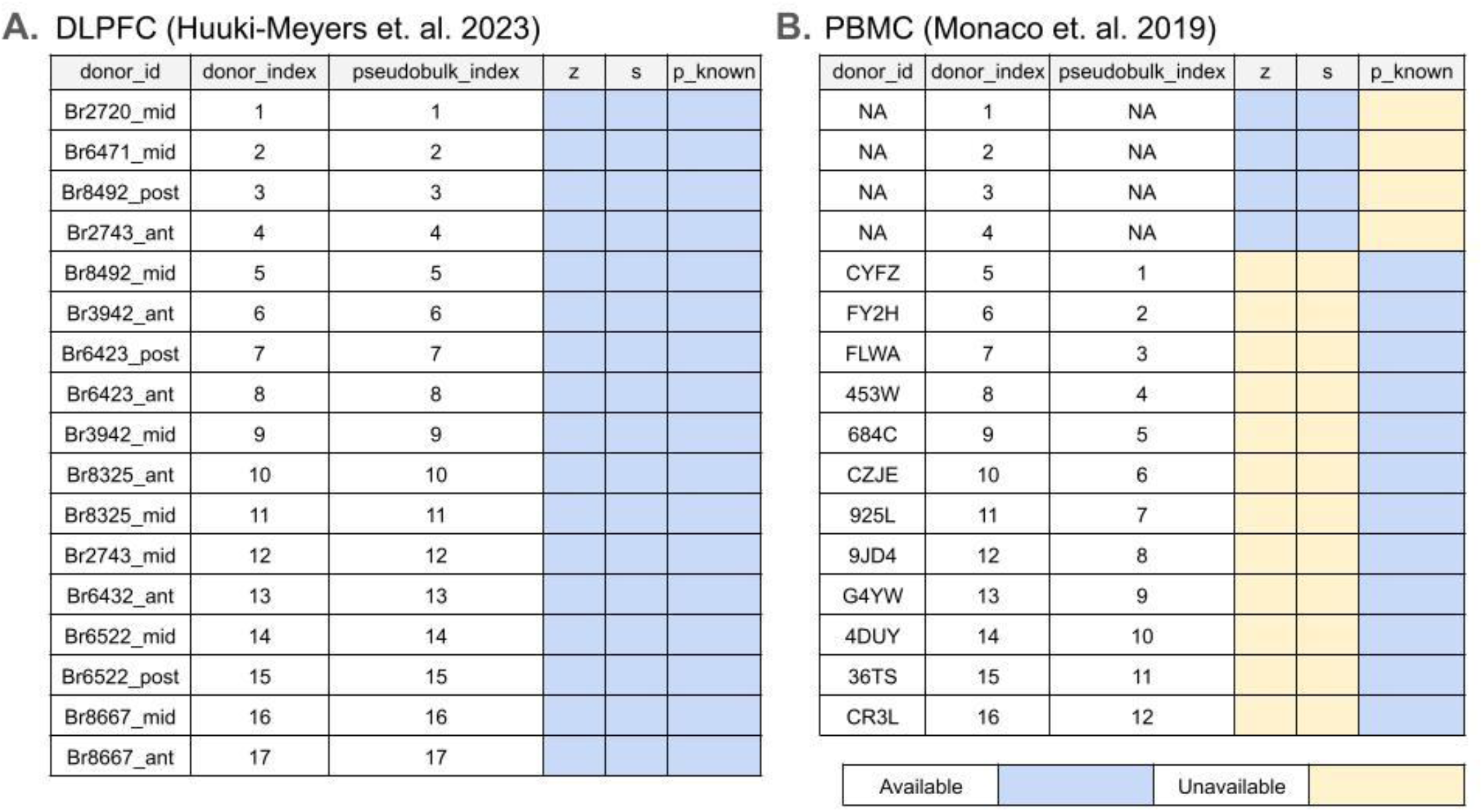
Availability of samples by cohort for pseudobulk experiments in two cohorts and two tissues. (**A**) Availability of samples for pseudobulk experiments in Huuki-Myers et al. (46), with columns indicating (left to right) donor identifier, donor index, pseudobulk index, *Z, S*, and *P*_*known*. (**B**) Availability of samples for pseudobulk experiments in Monaco et. al. 2019 (50), with columns indicating (left to right) donor identifier, donor index, pseudobulk index, *Z, S*, and *P*_*known*. Details about pseudobulk data types provided in **Methods**. Cell colors for *Z, S*, and *P*_*known* indicate data was either (blue) available or (yellow) unavailable for analysis.

## Supplemental tables

**Table S1** | **Tissue block-level summary of Huuki-Myers et al. (2023) (46) DLPFC *N*=17 snRNA-seq libraries generated from tissue blocks obtained from 10 adult neurotypical donors across three position across the dorsolateral prefrontal cortex (DLPFC)**. These snRNA-seq samples were used in the *in silico* pseudobulk experiments in this paper. Columns summarize information about the snRNA-seq libraries including the number of donors the tissue blocks originated from, the percent of samples that are female, DLPFC anterior to posterior position (position), percent of DLPFC samples stratified by position (posterior, middle, anterior), and the number of nuclei per sample (mean, median, sd, total).

**Table S2** | **Cell type-level summary of Huuki-Myers et al. (2023) (46) DLPFC *N*=17 snRNA-seq libraries generated from tissue blocks obtained from 10 adult neurotypical donors across three regions of the dorsolateral prefrontal cortex (DLPFC)**. Summaries are aggregated to *k*=2 (first two rows) or *k*=3 (last two rows) cell types. Columns include *k* dimensions (total cell types), cell type label, nuclei summaries by tissue block (median, mean, sd), proportion summaries (median, mean, sd), library summaries (mean, sd), and pseudobulk cell type scale factor.

**Table S3** | **Results from estimating the cell composition using the pseudobulk tissue samples from three different sources of snRNA-seq**. Three sources of data include snRNA-seq libraries from Huuki-Myers et al. 2023, Tran et al. 2021, and Monaco et al. 2019. Columns include condition (“withscale” if cell size adjustment was used, “noscale” if no adjustment was used, “all” if both adjustment conditions were combined, and an algorithm name where *NNLS* was used if no algorithm name was specified), root mean squared error (RMSE), dataset (represents the source of where the data came from), k_total (total cell types considered in pseudobulk experiment), k_rmse (total cell types used in RMSE calculation), cell type labels in the calculation (separated by “;”), and experiment (type of experiment performed, either “pseudobulk” where pseudobulks were tested, “shuffle” where cell sizes were shuffled across references, or “bulk” where real bulk RNA-seq samples were used).

**Table S4** | **Sample-level summary of Tran et al. (2021) (49) DLPFC *N*=3 snRNA-seq libraries generated from all posterior tissue blocks in DLPFC obtained from 3 adult neurotypical donors**. These snRNA-seq samples were used in the *in silico* pseudobulk experiments in this paper. Columns summarize information about the snRNA-seq libraries including the number of donors the tissue blocks originated from, the percent of samples that are female, DLPFC orientation (region), percent by DLPFC samples stratified by subregion (posterior, middle, anterior), and the number of nuclei per sample (mean, median, sd, total).

**Table S5** | **Cell type-level summary of Tran et al. (2021) (49) DLPFC *N*=3 snRNA-seq libraries generated from 3 adult neurotypical donors across the posterior region of the dorsolateral prefrontal cortex (DLPFC)**. Summaries are aggregated to *k*=2 (first two rows) or *k*=3 (last two rows) cell types. Columns include *k* dimensions (total cell types), cell type label, nuclei summaries by tissue block (median, mean, sd), proportion summaries (median, mean, sd), library summaries (mean, sd), and pseudobulk cell type scale factor.

**Table S6** | **Summary of cell sizes estimated from Huuki-Myers et al. 2024 (26)**. Columns include sample id, DLPFC region, subject identifier (corresponding to first part of sample id), library-based cell size scale factors and their ratios from snRNA-seq (“sn”) and RNAScope (“rn”) for neuron and glial.

**Table S7** | **Results of shuffle analyses of Huuki-Myers et al. (2023) (46) DLPFC *N*=13 snRNA-seq libraries for neuron and glial cell types**. (left to right) Columns include known proportions, predictions, cell type label, sample id (id of sample with cell sizes for shuffle experiment), error, index sample id (id of source for pseudobulk), shuffle term (location in deconvolution function), and sizes of neuron and glial cells (from pseudobulk source or index sample id), and panel indicating the corresponding to **Figure S3** panels. Rows 2 and 3 correspond to the concordant sample (sample id equals index sample id column, Br3942_mid) featured in shuffle experiment in **Figure S3B**, and rows 21 and 22 correspond to concordant sample in **Figure S3A** (Br2743_ant).

**Table S8** | **Platform-level data summaries of Overview of paired bulk-RNA-seq, snRNA-seq and smFISH datasets from adult neurotypical postmortem human DLPFC tissue blocks**. Columns include platform name and sample preparation, and quantities of samples and sample sources (“donors”).

## Notes

### Competing Interest Statement

The authors have declared no competing interest.

